# Sensitive protein detection using site-specifically oligonucleotide-conjugated nanobodies

**DOI:** 10.1101/2022.01.10.475658

**Authors:** Rasel A. Al-Amin, Phathutshedzo M. Muthelo, Eldar Abdurakhmanov, Cécile Vincke, Shahnaz P. Amin, Serge Muyldermans, U. Helena Danielson, Ulf Landegren

## Abstract

High-quality affinity probes are critical for sensitive and specific protein detection, in particular for detection of protein biomarkers at early phases of disease development. Proximity extension assays (PEA) have been used for high-throughput multiplexed protein detection of up to a few thousand different proteins in one or a few microliters of plasma. Clonal affinity reagents can offer advantages over the commonly used polyclonal antibodies (pAbs) in terms of reproducibility and standardization of such assays. Here we explore nanobodies (Nb) as an alternative to pAbs as affinity reagents for PEA. We describe an efficient site-specific approach for preparing high-quality oligo-conjugated Nb probes via enzyme coupling using Sortase A (SrtA). The procedure allows convenient removal of unconjugated affinity reagents after conjugation. The purified high-grade Nb probes were used in PEA and the reactions provided an efficient means to select optimal pairs of binding reagents from a group of affinity reagents. We demonstrate that Nb-based PEA (nano-PEA) for interleukin-6 (IL6) detection can augment assay performance, compared to the use of pAb probes. We identify and validate Nb combinations capable of binding in pairs without competition for IL6 antigen detection by PEA.

## INTRODUCTION

Progress in technologies for protein detection and analysis enables studies that go beyond investigations of genetic predisposition at the level of DNA. Most drugs act by interfering with protein function and proteins are therefore highly relevant as targets for analysis. Also, for diagnostic purposes protein analyses can provide insights in dynamic states by monitoring protein levels in samples collected at successive intervals. RNA analyses can also provide insights in activity states, but they typically require access to the tissues where the corresponding genes are expressed. Moreover, protein levels and activities often depend on protein degradation rates, post-translational modifications, and their involvement in forming complexes that profoundly affect their activities, neither of which can be predicted from the levels of the corresponding transcripts. Immunoassays are the most commonly used methods for clinical proteomic analysis, widely applied to diagnose disease and to monitor therapeutic effects clinically and during drug development.

The success of affinity-based immunoassays depends on the availability of a large repertoire of affinity reagents against most clinically relevant protein targets.^1^ In addition, molecular protein detection assays with improved proofreading are needed for efficient detection of target molecules at high efficiency and with minimal nonspecific background to ensure highly specific and sensitive detection. Oligo-assisted proximity-based immunoassays^2^ have been adapted for different proteomic applications, such as for high-throughput plasma proteomics,^3^ visualization of protein and their complexes *in situ*,^4^ detection of drug-target engagement and of extracellular vesicles,^5-6^ infectious diagnostics,^7^ flow cytometry^6^ and western blotting.^8^ Variants of proximity-based assays have been developed where each target molecule must be recognized by three different Abs, for enhanced detection in solution-phase assays^9^ or on solid supports^10^ and prostate derived microvesicles called prostasomes have been detected at elevated levels in plasma from prostate cancer patients using sets of five different Abs.^11^

The proximity extension assay (PEA) is a homogenous immunoassay using pairs of Abs conjugated to oligos with free, complementary 3’ ends.^3^ When a pair of Ab conjugates bind their target protein, the attached oligos are brought in proximity, permitting extension by a polymerase to produce a DNA reporter molecule for detection by real time PCR or sequencing.^3^ Around a hundred or even more different proteins can be measured in 1 μl sample aliquots using this technique, but access to suitable affinity reagents represents a limiting factor. Recombinant protein-binding reagents have the advantages over pAbs that they can be produced in any desired amount, and they can be engineered for site-directed conjugation of precisely one oligo per molecule, simplifying standardization.^12-14^ Examples of such affinity reagents to replace or complement Abs include nanobodies (Nbs), single-chain variable fragments (scFvs), monobodies and designed ankyrin repeat proteins (DARPins).^14^

Nbs are single-domain Ab fragments of 120-130 amino acids containing a single conserved disulphide bridge, and they are derived from the variable regions of atypical immunoglobulins of *Camelidae*.^15-16^ Nbs have proven useful as high-affinity reagents for research, diagnostics and therapeutics owing to their high specificity, small size (∼15 kDa) and straightforward bacterial expression.^15-18^ These minimal protein domains can exhibit high-affinity binding for protein targets, and they may be site-directedly modified using a convenient, recently described enzyme-based conjugation technique which relies on Staphylococcus aureus Sortase A (SrtA)- mediated coupling reactions.^19^

Here, a set of Nbs were modified with sortase tags and conjugated to suitably modified oligos by SrtA-coupling. These four different IL6 sortase-tag Nb clones (NbSORIL6) reagents were explored as substitutes for or complements to pAbs in PEA. We established assays combining pairs of Nb reagents, and compared the efficiency of protein detection by the different pairs of reagents. The Nb reagents allowed sensitive protein detection by PEA with real-time qPCR readout.

## EXPERIMENTAL SECTION

### Surface plasmon resonance (SPR) biosensor analysis

The affinity of the six SORIL6 Nbs for human IL6 was determined by SPR, using the BIACORE-T200 (Cytiva, Freiburg, Germany). The experiments were performed by direct immobilization of the recombinant IL6 protein (Cat. No. Z03034, Genscript), on a CM5 biosensor chip surface (Cytiva) by a standard amine coupling procedure using NHS (N-hydroxysuccinimide)/EDC (1-ethyl-3-(3-dimethylaminopropyl)carbodiimide hydrochloride) chemistry, resulting in a final change of 200 Response Units (RU). The formula used to calculate how much immobilized IL6 to achieve: [Rmax (200RU) = immobilized IL6 (200 (Rmax) x 21,041 (MW IL6)/ 15000 (MW Nb)]. The SPR measurements were performed at 25°C with HBS (10 mM of HEPES pH 7.4, 150 mM of NaCl, 0.005% Tween-20, and 3.4 mM EDTA) as running buffer. For affinity measurements, the purified Nbs were injected sequentially in 2-fold serial dilutions, from 500 to 1.95 nM, at a flow rate of 30 µl/min. The association step was followed for 120 s, the dissociation step was 600 s, and a regeneration of 60 s at 30 μl/min was performed using 100 mM glycine (pH 2.7) was performed. After the regeneration step, an additional stabilization step of 180 s was included. The kinetic rate constants were determined by global fitting of a 1:1 binding model with drift to the sensorgrams using the BIACORE Evaluation software (Cytiva, Freiburg, Germany), and the ratio between the rate constants (k_off_/k_on_) was used to determine the equilibrium dissociation constant (K_D_). For the epitope binning experiments, measurements were performed at 10 µl/min in HBS buffer. The first sample consisting of an excess of a first Nb (NbX), at a concentration of 200 times its K_D_ value, was initially injected for 300 s to saturate all its available epitopes on the IL6 protein. This step was followed by a second injection under the same conditions for 300 s, but with a mixture of e.g., NbX+NbY (both at a concentration of 200 times their respective K_D_ values, to guarantee equal competition of all Nbs). The RU was then monitored for 600 s, followed by a regeneration step of 90 s at 30 μl/min, using 100 mM of glycine (pH 2.7). The next cycle was initiated after a stabilization time of 180 s. All Nb-pairs were evaluated in all possible combinations, and for each pairwise combination 4 cycles were performed:

1. NbX + NbX
2. NbX + Nb(X and Y)
3. NbY + Nb(X and Y) and
4. NbY + NbY

### Nb-oligo conjugation, identification and purification

Hundred μL reactions were prepared with 24, 60 and 120 μM Nbs (2-, 5- and 10-fold molar excess of NbSORIL6 over oligos), 150 μM SrtA enzyme and 12 μM oligo modified at the 5’ end with three glycines (GGG-oligo) in SrtA coupling buffer (50 mM Tris pH 8.0, 150 mM NaCl, 10 mM CaCl_2_; pH7.5-8.0). The GGG-modified oligos F1 and R1 or H1 and H2 were conjugated at stoichiometric ratios to Nb and a 5-fold molar excess of Nb over oligos were found to be optimal. The reactions were incubated at 37°C overnight with gentle shaking on a rotating platform. Following incubation, the remaining His-tagged SrtA enzyme, acyl-intermediate, unconjugated Nb and cleaved-off his tags were removed by binding to Dynabeads® Magnetic Beads (ThermoScientific) overnight at 4°C with continuous shaking in binding buffer (100 mM sodium-phosphate, pH 8.0, 600 mM NaCl, 0.02% Tween®-20). Conjugates were validated on 4-12% NuPAGE Bis-Tris Gel (ThermoScientific) with NuPAGE™ LDS Sample Buffer (4X) (ThermoScientific) loading sample buffer + 50 mM DTT heated at 80°C for 3 min, followed by electrophoresis in 1x MES-SDS buffer for 60 min at room temperature. The conjugates were visualized by staining using the PlusOne DNA Silver Staining Kit (GE Healthcare, Uppsala) and Gel Doc XR+ (Bio-Rad) for imaging (Figure 2A and Supplementary Figures S6). To ensure thorough purification, the purification process was repeated up to three times on new beads. The conjugates were purified from unreacted oligos and any remaining free Nb by HPLC using a proFIRE instrument (Dynamic Biosensors, Planegg, Germany) according to the manufacturer’s instructions. In brief, the ion exchange column was pre-equilibrated with buffer A (50 mM Na_2_HPO_4_/NaH_2_PO_4_ pH 7.2, 150 mM NaCl) followed by injection of 160 µl of beads purified Nb-oligo conjugates. The conjugates were eluted by a pre-defined salt gradient using buffer B (50 mM Na_2_HPO_4_/NaH_2_PO_4_ pH 7.2, 1 M NaCl) at a flow rate of 1 ml/min. Subsequently, the PEA assay-specific 89-mer oligo HF1, 73-mer oligo HR1 or 111-mer oligo DimerS1-F, 121-mer oligo DimerS1-R or 121-mer oligo DimerS2-F, 119-mer oligo Dimer-S2 comprising 37-nt and 9-nt segments complementary to oligos F1 and R1 were mixed together to prepare a hybrid.

### Validation of Nb-oligo probes and comparison via SPR

The binding of 4 different Nbs, and Nb-oligo conjugates of two of them, to IL6 was investigated using two approaches: (i) IL6 was directly immobilized on a CM5 biosensor chip surface (GE Healthcare, Uppsala, Sweden) by a standard amine coupling, resulting in immobilization of 700-900 RU in 10 mM sodium acetate buffer, pH 4.5. The Nbs were injected over the IL6-coated chip surface in a series of concentrations from 15.6 nM to 250 nM in the running buffer of 20 mM Tris-HCl pH 8.0, 125 mM NaCl, 0.05% Tween-20, at a flow rate of 30 µl/min. (ii) IL6 was captured to a level of 1300-2800 RU on an anti-IL6 monoclonal antibody (mAb clone 13A5, MABTECH) coupled to the chip surface and the interaction analysis of 4 selected Nbs was performed similarly as described above. Anti-IL6 mAb was immobilized on the chip surface by amine coupling in 10 mM sodium acetate buffer (pH 5.0), resulting in an immobilization level of 15000-19000 RU. Reference surface and blank were subtracted from the sensorgrams. The data were analyzed using Biacore T200 Evaluation software v. 3.0 (GE Healthcare, Uppsala, Sweden).

### Preparation of PEA Ab probes

Anti-human IL6 pAbs were diluted in PBS at 2 µg/µl and stored at -20°C. Anti-human IL6 pAbs (2 µg/µl in PBS) were activated with a 20-fold molar excess of the cross linker dibenzylcyclooctyne-NHS (DBCO-NHS) ester (CLK-A102N, Jena Bioscience), diluted in dimethyl sulfoxide (DMSO) at 4 mM, and incubated at RT for 30 min. The reaction was stopped by adding of 100 mM Tris-HCl, pH 8.0 and incubating at RT for 5 min. The excess unreacted DBCO-NHS ester was removed from the activated Ab with an equilibrated Zeba Spin Desalting Column (7k MWCO, Thermo Scientific). After purification, the DBCO-labeled Abs were mixed with a 2.5-fold molar excess of 5’ azide-modified forward oligo F1 or reverse oligo R1 and incubated overnight at 4°C. Anti-human IL6 Ab concentrations were quantified using the Quant-it protein assay kit (Life Technologies). Ab conjugates were validated on 10% TBE urea denaturing gels and validated on an agarose gel (Supplementary Figure S8). Then, an assay-specific 89-mer oligo HF1 and 73-mer oligo HR1 comprising 37-nt and 9-nt segments complementary to oligos F1 and R1, were combined to prepare DNA hybrids with the pAb conjugated oligos F1 or R1 (Supplementary Figure S1). In addition, goat IgG was conjugated with both GGG-modified oligos F1 or R1 and used as extension and PCR control. The PEA probes were stored at 250 nM in PBS with 0.1% BSA and 0.05% NaN_3_ and the probes were diluted to a 1.33 nM stock concentration in PEA buffer before use.

### PEA reactions and data analysis

Pairs of oligo-conjugated Nb or anti-human IL6 Ab probes were mixed in assay diluent at a final concentration of 100 pM each, and 3 μl of the probe mixture was added per microtiter well, followed by addition of 2 μl IL6 dilutions. Recombinant human IL6 protein had been diluted in assay diluent (PBS with 5 mM EDTA, 100 μg/ml single-stranded salmon sperm DNA (Sigma Aldrich), 0.1% BSA, 1 mM biotin, 100 nM goat IgG, 0.05% Tween20 solution). After incubation at 37°C for 1 hour, 46 μl extension solution was added, containing 1X Hypernova buffer (BLIRT S.A.), 1.5 mM MgCl_2_, 0.2 mM of each dNTP, 1 μM of each FEP (forward extension primer) and REP (reverse extension primer), 0.2 U/ml DNA Polymerase (Invitrogen) and 1 U/ml Hypernova DNA polymerase. The extension reactions were conducted at 50°C for 20 min, followed by a 5-min heat-activation step at 95°C and 17 cycles of pre-PCR of 30 sec at 95°C, 1 min at 54° C, and 1 min at 60° C. For the subsequent qPCR detection, 2.5 μl of extension/pre-PCR products were transferred to a 96 or 384-well plate and combined with 7.5 μl qPCR mix containing 1X PCR buffer (Invitrogen), 0.1 µM of each FEP (forward extension primer) and REP (reverse extension primer) PCR primers, 2.5 mM MgC1_2_, 0.2 µM TaqMan probe (or 0.5x SYBER green I), 0.25 mM dNTPs (including dUTP instead of dTTP), 0.02 U/µl Klenow exo-, 0.02 U/µl uracil-N-glycosylase, 1.5 U/µl Taq polymerase and 1.33 μM ROX (ROXTTTTTTT, Biomers). Quantitative real-time PCR was performed with an initial incubation at 25°C for 30 min, denaturation at 95°C for 5 min, followed by 40 cycles of 15 sec denaturation at 95°C, and 1 min annealing/extension at 60°C. ROX was used as reference fluorescence and SYBR green as detection fluorescence. The qPCR data were recorded as Ct (cycle threshold) values. The data were analyzed with Microsoft Excel. The plots were generated on Microsoft Excel and using an in-house script developed in ‘Rstudio’ (http://www.R-project.org/). The Ct values were plotted along the y-axis against the concentration of target protein in the reactions (x-axis). The data was analyzed by linear regression to determine the LOD (limit of detection). The LOD was defined as the concentration of protein corresponding to Ct_LOD_=Ct_N_ − (3 x S_N_), where Ct_N_ is the background noise corresponding to the mean value of the three negative control samples (N), and S_N_ is the mean standard deviation of those values.

## RESULTS AND DISCUSSION

### Site-specific conjugation of oligos to NbSORIL6, validation and two-step purification of conjugates

Typically, conventional sandwich immunoassays require chemical modification of the Abs in order to attach reporter groups, but conjugation to random sites risk compromizing target binding. In our model experiments we used four recombinant Nbs directed against the IL6 protein, site-specifically conjugating exactly one oligo per Nb molecule by SrtA coupling (Figure 1, 2A and Supplementary Figure S6). The SrtA enzyme reacts with the consensus sequence LPETGG of the Nb and cleaves the peptide bond between threonine and glycine, subsequently joining the threonine by a peptide bond to the N-terminal glycine on the oligo to give rise to the sequence LPETGGG (Figure 1).^19, 20^ This site-directed conjugation technique has been previously shown to allow construction and purification of well-defined protein conjugates. Conjugation of NbSORIL6 to the glycine-oligos results in removal of the His-tag. Any remaining unconjugated Nbs as well as the His-tagged SrtA were conveniently removed through incubation with anti-his magnetic Ni-beads (Figure 1). The pure Nb-oligos conjugates were then isolated from unreacted oligos and excess free Nbs by chromatography using a proFIRE instrument. The Nb conjugates were well separated from free Nb and oligos yielding 68% and 87% pure conjugates for NbSORIL6_15-oligos H1 and NbSORIL6_16-oligos H2 respectively, and the yield was calculated by comparing the material used for the conjugation reaction and the recovered material after chromatography and elution (Figure 2B-D). Each Nb was conjugated to either of two oligos to which secondary oligos were then hybridized to create reagents for PEA reactions (Supplementary Figure S1). Polyclonal Ab-based PEA probes were prepared through click chemical conjugation in a non-site-specific manner. The oligos are coupled to primary amines on the Ab via heterobifunctional crosslinkers. The Ab conjugates were validated by agarose gel analysis, showing the presence of variable numbers of oligos per Ab (Supplementary Figure S8).

**Figure 1.**
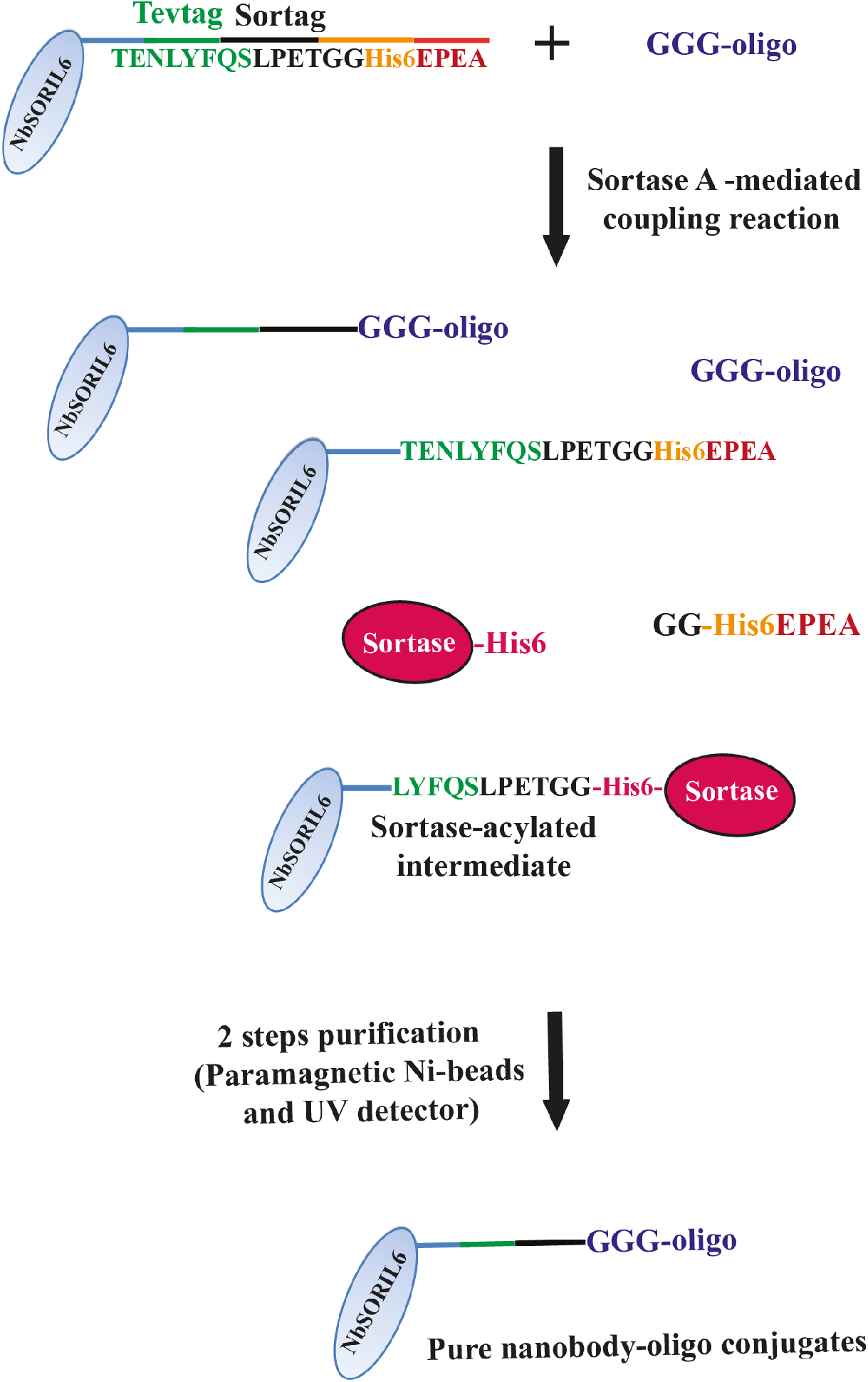
Schematic illustration of the site-specific conjugation of PEA oligos to the Nbs. The Nbs were expressed with a sortase tag sequence (LPETGG) recognized by SrtA enzyme, followed at the end by a His-tag at the C-terminus, and the oligos to be conjugated had three glycines at their 5’ ends as required for joining to the Nb by SrtA. The SrtA enzyme also included a sacrificial His-tag at the C-terminus, which allowed for removal of any unconjugated reactants remaining in the solution by reverse nickel affinity pull down with paramagnetic Ni-beads. After the SrtA reaction all His-tagged material in the solution was collected on paramagnetic Ni-beads and discarded, while conjugates and free oligos were obtained in solution. This was followed by separation of the conjugated oligos using a proFIRE instrument (Dynamic Biosensors) monitored via UV detection to isolate pure conjugates.

**Figure 2.**
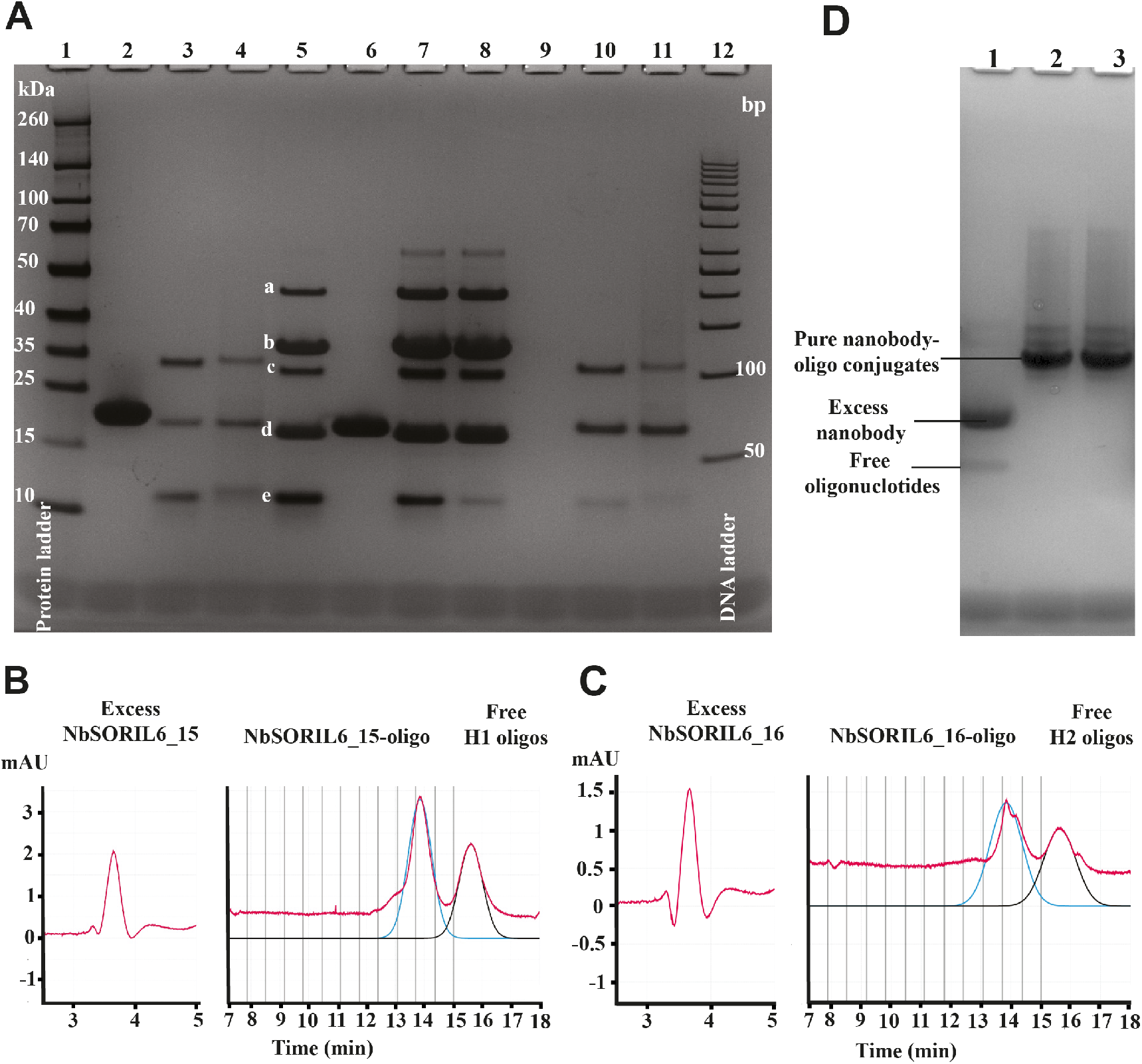
Gel validation of oligo-Nb conjugation reaction, two-step purification and validation of the pure NbSORIL6-oligo conjugates. **(A)** Gel electrophoresis of sortase-mediated oligo-Nb conjugation reaction products and samples undergoing two-step purification. Lane 1: protein ladder, lane 12: gene ruler 50 bp DNA ladder, lanes 2 and 6: free NbSORIL6_16 and NbSORIL6_15 respectively, lane 9: negative control (only loading buffer). Unpurified reaction products with a 2, 5 or 10-fold molar excess of NbSORIL6_15 over oligo H1 is seen in lanes 5, 7 and 8, respectively. The species identified in order of increasing migration in lane 5 are a) acylated product, b) free SrtA enzyme, c) Nb-oligo conjugate, d) excess unreacted NbSORIL6_15, and e) free oligo H1. Lanes 3 and 4 show the products NbSORIL6_16-oligo H1 and NbSORIL6_16-oligo H2, respectively, after a single incubation with Ni-beads, while lanes 10 and 11 demonstrate products from NbSORIL6_16-oligo H1 and NbSORIL6_16-oligo H2 reactions, respectively remaining in solution after two incubations with the beads. **(B-C)** The products shown in lanes 10 and 11 of the gel were separated from free Nb and unreacted oligos using the proFIRE instrument, yielding the pure Nb conjugates NbSORIL6_15 **(B)** and NbSORIL6_16 **(C)**. The trace of the separation is shown in red. The software estimates the amount of conjugates and unreacted oligos by fitting the chromatogram peaks. The blue lines correspond to fitted peaks of conjugates, while black lines represent fitted peaks of unreacted oligos. **(D)** Gel electrophoretic validation of sortase-mediated oligo-Nb conjugation reaction products after two-step purification. Lane 1: free NbSORIL6_16 and free oligo H2. Lanes 2 and 3 demonstrate products from NbSORIL6_16-oligo H1 and NbSORIL6_16-oligo H2 reactions, respectively. The products shown in lanes 2 and 3 of the gel were separated from free Nb and unreacted oligos using the proFIRE instrument, yielding the pure Nb conjugates.

### Characterization of binding affinity using SPR biosensor analysis

The interaction between recombinant human IL6 antigen and the 4 different NbSORIL6 reagents as well as the NbSORIL6_21 and _15 oligo conjugates were validated by SPR biosensor analysis (Table S1 and Supplementary Figures S2-3). The NbSORIL6_21 and _15 and their respective conjugates all showed an interaction with the immobilized IL6. Some of the interactions were well described by a simple a 1:1 interaction model, allowing the affinities to be estimated from a global analysis of the complete sensorgrams: NbSORIL6_5 (K_D_ = 6.9 × 10^−10^ M), NbSORIL6_15 (K_D_ = 1.1 × 10^−8^ M), NbSORIL6_21 (K_D_ = 7.4 × 10^−9^ M), NbSORIL6_21-oligo H2 conjugates (K_D_ = 4.4 × 10^−8^ M) and NbSORIL6_16 (K_D_ = 4.7 × 10^−9^ M). A steady-state analysis of the sensorgrams was performed as a control. The data showed that of the different NbSORIL6 variants tested, NbSORIL6_15 had the weakest affinity for IL6 (Table 1 and Supplementary Tables S2, 4). The NbSORIL6_21-oligo conjugate showed a different kinetic profile, with a slower association and dissociation rates compared to the unconjugated NbSORIL6_21 (Supplementary Figures 3-4).

**Table 1.** Equilibrium dissociation constants obtained from different interaction models between IL6, immobilized on the chip surface, and Nbs. The corresponding equilibrium dissociation constants (*K*_D_) were determined by various models are presented in the table below:

| Nbs | $K_D$ (M) 1:1 | $K_D$ (M) Steady-state |
| --- | --- | --- |
| NbSORIL6_5 | $6.9 \times 10^{-10}$ | $1.6 \times 10^{-10}$ |
| NbSORIL6_15 | $1.1 \times 10^{-8}$ | $3 \times 10^{-8}$ |
| NbSORIL6_16 | $4.7 \times 10^{-9}$ | $1.1 \times 10^{-8}$ |
| NbSORIL6_21 | $7.4 \times 10^{-9}$ | $2.1 \times 10^{-8}$ |
| NbSORIL6_21-oligo conjugates | $4.4 \times 10^{-8}$ | Not determined |

We included a mAb specific for human IL6-specific as a positive control in our analysis, which showed that the mAb binds much stronger than all Nbs tested with almost no observable dissociation rate from the target IL6 protein, preventing an accurate quantification of its affinity (Supplementary Figure S3). It is however to be expected that bivalent mAbs via avidity bind with higher (apparent) affinity via avidity effects, compared to the monovalent Nbs. We further investigated the affinities of all four NbSORIL6s to IL6 captured to the surface via a mAb immobilized on the sensor surface. Under these conditions, the binding of NbSORIL6_15 and _21 was significantly reduced (Supplementary Figure S5 and Supplementary Table S4), probably due to competition for the binding site with this mAb. However, the affinity of NbSORIL6_5 was only slightly reduced (Table 1 and Supplementary Table S4). As for NbSORIL6_16, the affinity as well as the binding level increased significantly when IL6 antigen was captured via an immobilized mAb rather than via direct immobilization. This could be due to improved orientation or epitope exposure of IL6 for binding by NbSORIL6_16 (Table 1 and Supplementary Table S4).

### Identifying non-overlapping binding pairs via PEA

The NbSORIL6-oligo conjugates were then explored for their potential to expand the range of reagents used for PEA, complementing Abs. Two aliquots of all NbSORIL6s were conjugated using SrtA to either of two oligos to which secondary oligos were then hybridized to create reagents for PEA reactions (Supplementary Figure S1A). We combined pairs of Nb probes at 100 pM with IL6 protein ranging from 100 ng/ml to 100 fg/ml and no added protein in PEA reactions (Figure 3B). We investigated three combinations of the reagents to see which would yield the highest sensitivity to detect target protein in PEA reactions. Of the evaluated pairs, the combination of NbSORIL6_5 and NbSORIL6_16 conjugates yielded the assay with the greatest sensitivity and a detection limit below 1 pg/ml (Figures 3B and 4A). We performed SPR epitope binning and compared all different pairwise combinations of SORIL6 Nbs from the set investigated herein (data not shown). Our identification of the best optimal Nb probe pair agreed with epitope mapping SPR data that showed that these two Nbs (SORIL6_5 and SORIL6_16) had the most effective binding as well as slower dissociation rates compared to the other combinations (Figure 4A-B). We did not detect any signal over background for this combination of two Nbs (SORIL6_5 and SORIL6_16) when the IL6 antigen was absent. Detection of IL6 using pAbs generated significantly higher signals, and detected lower concentrations of proteins could be detected compared to when Nbs PEA probes were used (Figure 3B). The much higher background for nanobodies, presumably because Nbs PEA probes were used at higher concentrations, or alternatively had a tendency to interact with each other.

**Figure 3.**
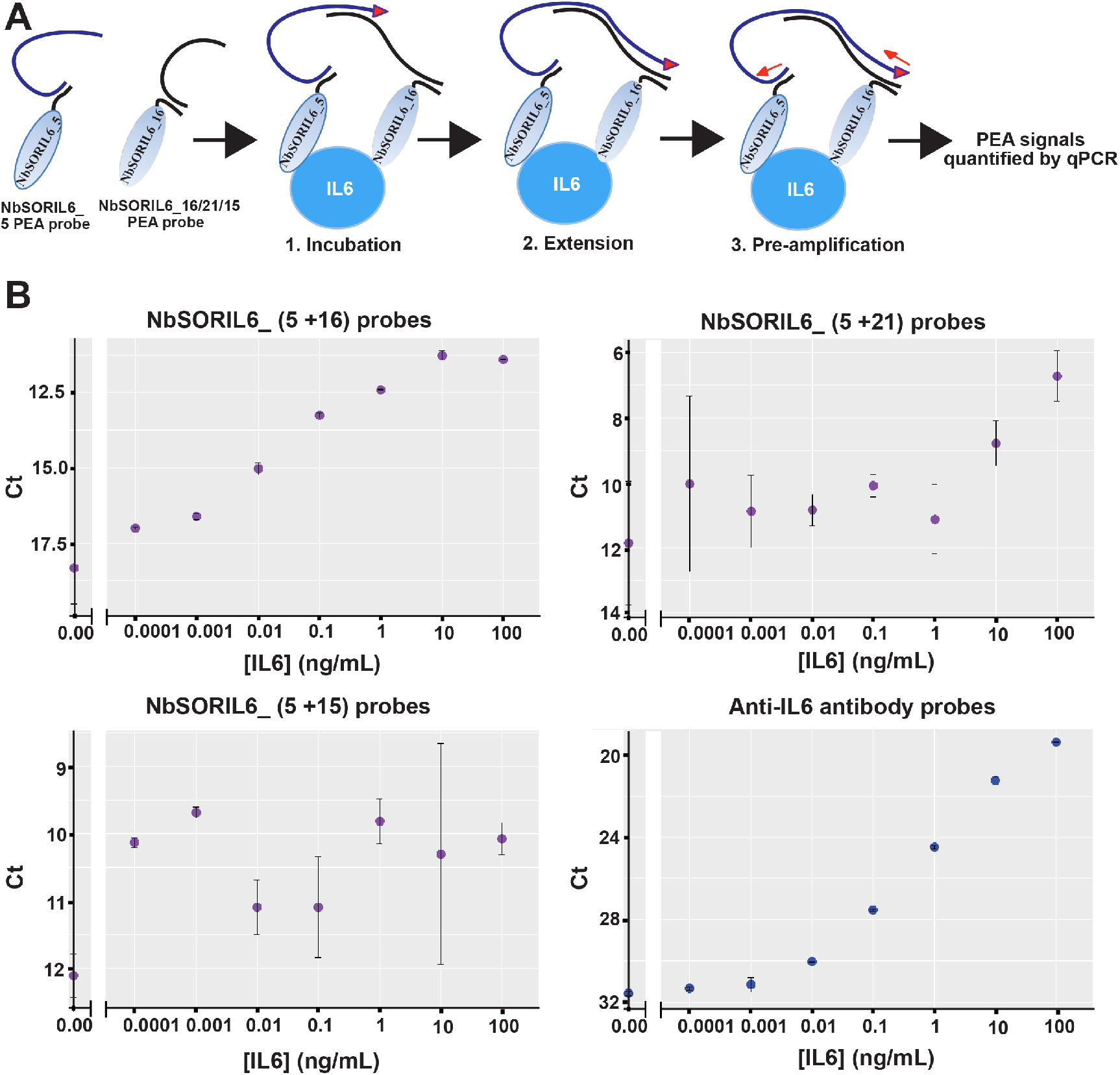
Nb-based detection of IL6 by PEA using Nb-oligo conjugates and regular pAb PEA probes. **(A)** Schematic illustration of nano-PEA reactions for protein detection. Nb probes pairs were incubated with target IL6 proteins, bringing the attached oligos in close proximity so that they can be mutually extended upon dilution and addition of a DNA polymerase. The resulting reporter DNA strands were quantified by real-time PCR (qPCR) as a measure of the amount of IL6 antigen in the buffer. **(B)** Different Nb probes combinations were applied for detection of serial dilutions of IL6 from 100 ng/ml to 0 fg/ml in PEA buffer. A pair of pAb-based PEA probes served as a positive control (PC) in a PEA detection. The data were analyzed using an in-house script developed in ‘Rstudio’ and the Ct values were plotted along the y-axis against the concentration (x-axis).

**Figure 4.**
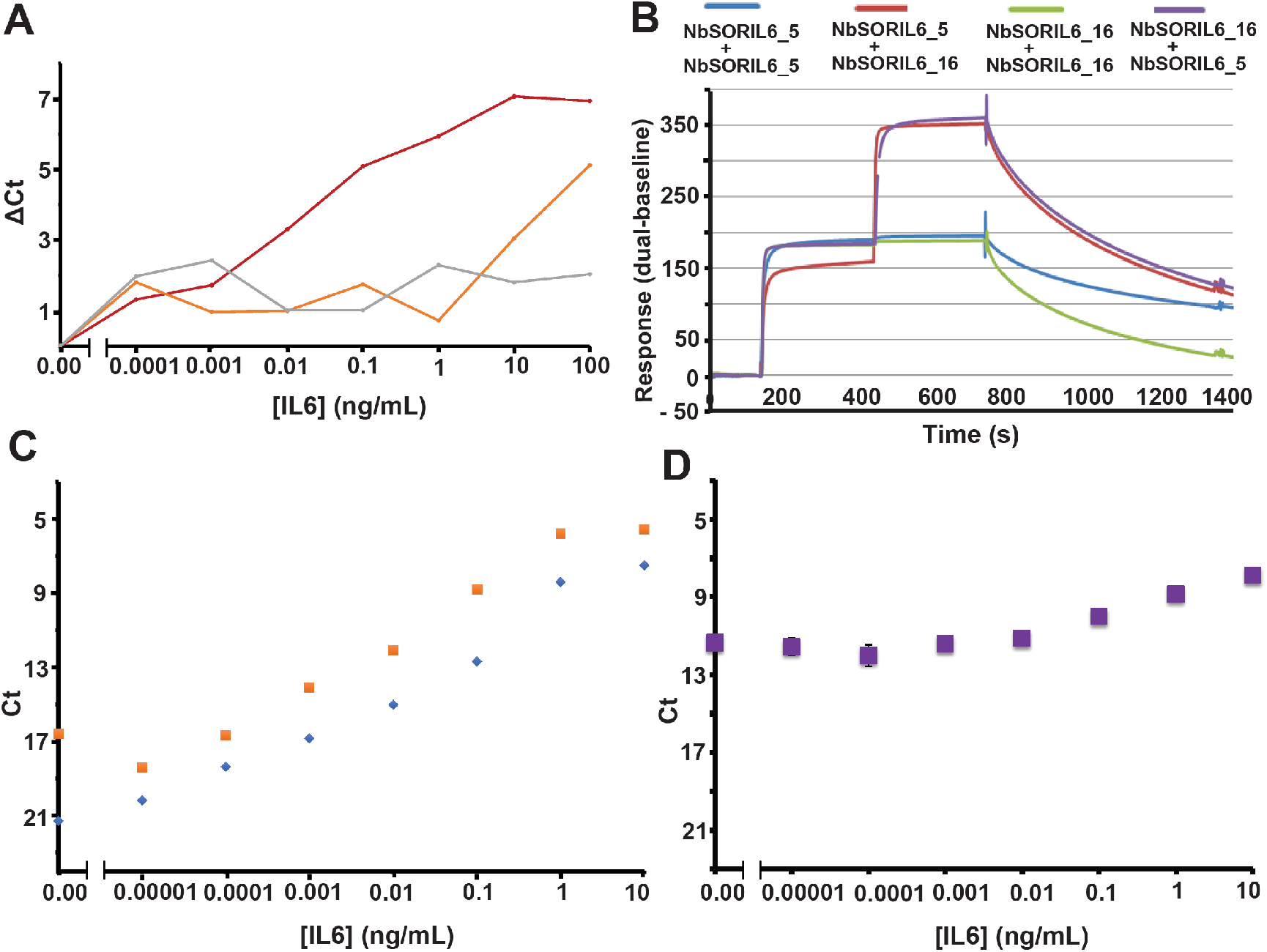
Comparison between Ab and Nb reagents for PEA. Detection of IL6 by PEA using NbSORIL6 pairs, pAb pairs or a combination of pAb and Nb probes. **(A)** Comparison of PEA results using NbSORIL6_5+ NbSORIL6_16 (dark red), NbSORIL6_5+ NbSORIL6_21 (orange) and NbSORIL6_5+ NbSORIL6_15 (gray). The x-axes show the concentration of IL6 antigen, while the y-axes show the cycle threshold values with background subtracted (delta Ct). **(B)** Epitope binning of NbSORIL6 via SPR with IL6 directly immobilized on the chip surface. Comparison of four pairwise combination of two Nbs (NbSORIL6_5 and NbSORIL6_16). Sensorgrams representing homo or hetero combinations interactions between NbSORIL6_5 and NbSORIL6_16. **(C)** Assessment of IL6 detection using either NbSORIL6_5 and NbSORIL6_16 (blue) or pAb pairs (orange) as PEA probes. The y-axes show the cycle threshold values (Ct) while the x-axes show the concentration of IL6 antigen. **(D)** Detection of IL6 antigen using a combination of NbSORIL6_5 probe and pAb.

The limit of detection (LOD) was calculated as described under data analysis in the Materials and Methods section and is represented in Supporting File 1. The optimal Nb PEA probes reached a plateau at the highest IL6 concentration, while the pAb PEA probes did not reach a plateau even at 100 ng/ml of IL6 (Figures 3B and 4C). Nb PEA reactions combining oligo-conjugated NbSORIL6_5 and _16 Nb pair detected lower concentrations of IL6 to the pAb PEA probes, and with a broader dynamic range (Figure 4C). PEA reactions combining NbSORIL6 probes 5 and 16 resulted in a LOD of 0.38 fM or 10 fg/ml, while PEA using pAb probes yielded a LOD of 3.2 fM or 80 fg/ml, indicating that the performance of NbSORIL6 probes can be comparable to that of pAb probes (Figure 4C and Table 2). To determine whether Nbs probes can be combined with Ab probes we performed PEA with a combination of Nb (NbSORIL6_5) and pAb PEA probes, however this assay performed poorly (Figure 4D). This could be because the probes have overlapping epitopes and therefore compete for binding.

**Table 2.**
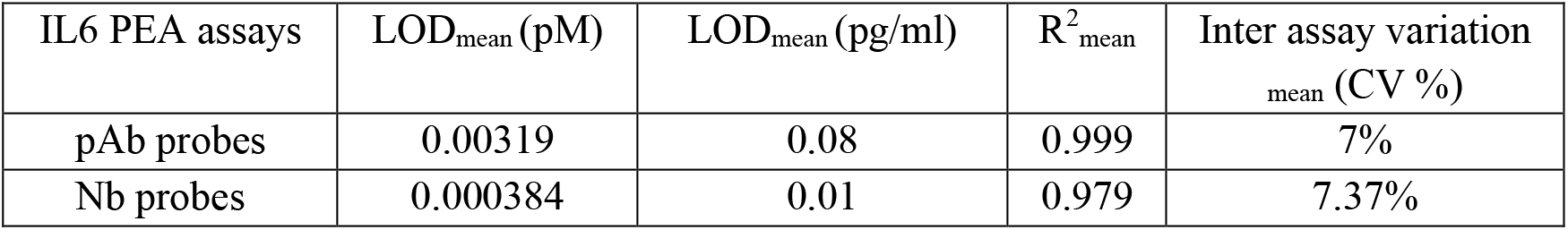
Analytical characteristic of pAb-based and nano-PEA assays, values are means of three separate experiments.

| IL6 PEA assays | LOD<sub>mean</sub> (pM) | LOD<sub>mean</sub> (pg/ml) | R<sup>2</sup><sub>mean</sub> | Inter assay variation<br>mean (CV %) |
| --- | --- | --- | --- | --- |
| pAb probes | 0.00319 | 0.08 | 0.999 | 7% |
| Nb probes | 0.000384 | 0.01 | 0.979 | 7.37% |

### Dimerization of Nb reagents

Dimerization of affinity reagents via their attached oligos could be expected to improve binding avidity and thus assay performance. We designed and evaluated the possibility to dimerize NbSORIL6-oligo conjugates in the PEA experiments using the best performing combination of NbSORIL6_5 and NbSORIL6_16 probes (Supplementary Figure S7A). We prepared homodimers of the conjugates through complementary oligos hybridization (Supplementary Table S3 and Supplementary Figure S1) to form homodimeric Nb PEA probes and the dimeric reagents were validated by gel electrophoresis (Supplementary Figure S7B). We established assays combining pairs of reagents and then compared the performance of the homo-dimers to their monomeric counterparts. However, under the investigated conditions no improved performance was observed using of homodimers of NbSORIL6_5+5 and NbSORIL6_16+16 conjugates over single Nbs combination of NbSORIL6_5 and NbSORIL6_16 (Supplementary Figure S7C).

## CONCLUSIONS

Abs randomly modified with DNA by chemical coupling can yield inconsistent results, affecting assay reproducibility.^21^ The site-directed attachment of exactly one oligo per Nb opens interesting possibilities where binders against different epitopes on the same protein molecule can be brought together by hybridization between their attached oligos for proximity ligation (PLA) or extension (PEA) assays. In this manner both efficiency and specificity of binding might be enhanced.^22-23^ Immunoassays that require dual recognition by binders provide higher specificity and can reduce the background compared to single binder-assays by reducing risks of cross reactivity by the affinity reagents for irrelevant target molecules.^24^ However, identifying non-competing pairs of binders remains a challenging task. Here we used PEA to identify optimized non-overlapping nanobody binding pairs. We compared different combinations of Nb pairs from 4 selected Nbs and we identified a pair that provided assay performance comparable to that of pAbs. The nano-PEA assay is a simple assay to establish that may be promising for future diagnostic multiplex assays that require highly specific and sensitive detection of sets of targets over wide ranges of protein concentrations.^25^

## ASSOCIATED CONTENT

### Supporting Information

The Supporting Information is available free of charge on the ACS Publications website:

Supporting information and results include methods, tables and figures; the experimental section and methods, expression, isolation of IL6-specific Nbs, affinity measurement of NbIL6s clones’ selection via ELISA and SPR, design of oligos for PEA probes, affinity validation of NbSORIL6s with or without conjugated oligos, gel electrophoresis validation of unpurified sortase-mediated conjugation reactions of four different Nbs, Comparison of the effect of dimerizing Nb probes, gel validation of Ab-oligos conjugation by copper free click chemistry. Supporting File 1; PEA assays performance comparison LOD calculations (XLSX).

## AUTHOR INFORMATION

### Author Contributions

R.A.A.-A., C.V., E.A. and P.M. performed most of the experimental work. C.V. produced and together S.M. validated the Nbs results. R.A.A.-A. conjugated with oligonucleotide, validated, purified of conjugates; E.A. and S.P.A. helped to analysis of validation and purifications results. C.V., E.A. and R.A.A.-A. performed SPR validation and H.D. helped to analysis data. P.M., R.A.A.-A., and U.L. designed and analyzed the PEA experiment. R.A.A.-A. wrote the manuscript and all authors contributed to the final paper.

### Notes

U.L. is a co-founder and shareholder of Olink Proteomics, having rights to the PEA technology.

## ACKNOWLEDGEMENTS

We thank Dr. Junhong Yan for discussions about Nbs oligo conjugation, and the Single Cell Proteomic Facility at SciLifeLab, Uppsala for support establishing PEA assays. Swedish Research Council’s [2012-5852, 2020-02258 to U.L.]; The Swedish Foundation for Strategic Research (SB16-0046); Torsten Söderbergs Stiftelse [M130/16 to U.L.]; The European Research Council under the European Union’s Seventh Framework Programme [FP/2007-2013/ ERC Grant Agreement No. 294409 to U.L.]; The Swedish Collegium for Advanced Studies (SCAS).

## For Table of Contents only

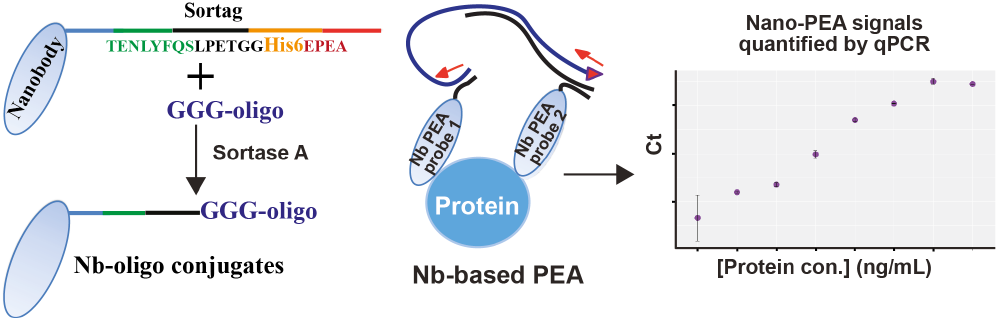

## REFERENCES

(1) Colwill K; Renewable Protein Binder Working Group, Gräslund S. A roadmap to generate renewable protein binders to the human proteome. Nat. Methods 2011, 8, 551–558.

(2) Fredriksson, S.; Gullberg, M.; Jarvius, J.; Olsson, C.; Pietras, K.; Gústafsdóttir, S. M.; Ostman, A.; Landegren, U. Protein detection using proximity-dependent DNA ligation assays. Nat. Biotechnol. 2002, 20, 473–477.

(3) Lundberg, M.; Eriksson, A.; Tran, B.; Assarsson, E.; Fredriksson, S. Homogeneous antibody-based proximity extension assays provide sensitive and specific detection of low-abundant proteins in human blood. Nucleic Acids Res. 2011, 39, e102.

(4) Söderberg, O.; Gullberg, M.; Jarvius, M.; Ridderstråle, K.; Leuchowius, K. J.; Jarvius, J.; Wester, K.; Hydbring, P.; Bahram, F.; Larsson, L. G.; Landegren, U. Direct observation of individual endogenous protein complexes in situ by proximity ligation. Nat. Methods 2006, 3, 995–1000.

(5) Al-Amin, R. A.; Gallant, C. J.; Muthelo, P. M.; Landegren, U. Sensitive measurement of drug-target engagement by a cellular thermal shift assay with multiplex proximity extension readout. Anal. Chem. 2021, 93, 10999–11009.

(6) Löf, L.; Ebai, T.; Dubois, L.; Wik, L.; Ronquist, K. G.; Nolander, O.; Lundin, E.; Söderberg, O.; Landegren, U.; Kamali-Moghaddam, M. Detecting individual extracellular vesicles using a multicolor in situ proximity ligation assay with flow cytometric readout. Sci. Rep. 2016, 6, 34358.

(7) Gustafsdottir, S. M.; Nordengrahn, A.; Fredriksson, S.; Wallgren, P.; Rivera, E.; Schallmeiner, E.; Merza, M.; Landegren, U. Detection of individual microbial pathogens by proximity ligation. Clin. Chem. 2006, 52, 1152–1160.

(8) Liu, Y.; Gu, J.; Hagner-McWhirter, Å.; Sathiyanarayanan, P.; Gullberg, M.; Söderberg, O.; Johansson, J.; Hammond, M.; Ivansson, D.; Landegren, U. Western blotting via proximity ligation for high performance protein analysis. Mol. Cell Proteomics 2011, 10, O111.011031.

(9) Schallmeiner, E.; Oksanen, E.; Ericsson, O.; Spångberg, L.; Eriksson, S.; Stenman, U. H.; Pettersson, K.; Landegren, U. Sensitive protein detection via triple-binder proximity ligation assays. Nat. Methods 2007, 4, 135–137.

(10) Darmanis, S.; Nong, R. Y.; Hammond, M.; Gu, J.; Alderborn, A.; Vänelid, J.; Siegbahn, A.; Gustafsdottir, S.; Ericsson, O.; Landegren, U.; Kamali-Moghaddam, M. Sensitive plasma protein analysis by microparticle-based proximity ligation assays. Mol. Cell Proteomics 2010, 9, 327–335.

(11) Tavoosidana, G.; Ronquist, G.; Darmanis, S.; Yan, J.; Carlsson, L.; Wu, D.; Conze, T.; Ek, P.; Semjonow, A.; Eltze, E.; Larsson, A.; Landegren, U. D.; Kamali-Moghaddam, M. Multiple recognition assay reveals prostasomes as promising plasma biomarkers for prostate cancer. Proc. Natl. Acad. Sci. U.S.A. 2011, 108, 8809–8814.

(12) Taussig, M. J.; Stoevesandt, O.; Borrebaeck, C. A.; Bradbury, A. R.; Cahill, D.; Cambillau, C.; de Daruvar, A.; Dübel, S.; Eichler, J.; Frank, R.; Gibson, T. J.; Gloriam, D.; Gold, L.; Herberg, F. W.; Hermjakob, H.; Hoheisel, J. D.; Joos, T. O.; Kallioniemi, O.; Koegl, M.; Konthur, Z.; … Uhlén, M. ProteomeBinders: planning a European resource of affinity reagents for analysis of the human proteome. Nat. Methods 2007, 4, 13–17.

(13) Baker, M. Reproducibility crisis: Blame it on the antibodies. Nature 2015, 521, 274–276.

(14) Harmansa, S.; Affolter, M. Protein binders and their applications in developmental biology. Development 2018, 145, dev148874.

(15) Helma, J.; Cardoso, M. C.; Muyldermans, S.; Leonhardt, H. Nanobodies and recombinant binders in cell biology. J. Cell Biol. 2015, 209, 633–644.

(16) Muyldermans S. Nanobodies: natural single-domain antibodies. Annu. Rev. Biochem. 2013, 82, 775–797.

(17) Vincke, C.; Gutiérrez, C.; Wernery, U.; Devoogdt, N.; Hassanzadeh-Ghassabeh, G.; Muyldermans, S. Generation of single domain antibody fragments derived from camelids and generation of manifold constructs. Methods Mol. Biol. 2012, 907, 145–176.

(18) Xiang, Y.; Nambulli, S.; Xiao, Z.; Liu, H.; Sang, Z.; Duprex, W. P.; Schneidman-Duhovny, D.; Zhang, C.; Shi, Y. Versatile and multivalent nanobodies efficiently neutralize SARS-CoV-2. Science 2020, 370, 1479–1484.

(19) Chen, L.; Cohen, J.; Song, X.; Zhao, A.; Ye, Z.; Feulner, C. J.; Doonan, P.; Somers, W.; Lin, L.; Chen, P. R. Improved variants of SrtA for site-specific conjugation on antibodies and proteins with high efficiency. Sci. Rep. 2016, 6, 31899.

(20) Lefranc M. P. IMGT, the international ImMunoGeneTics database. Nucleic Acids Res. 2003, 31, 307–310.

(21) Warden-Rothman, R.; Caturegli, I.; Popik, V.; Tsourkas, A.. Sortase-tag expressed protein ligation: combining protein purification and site-specific bioconjugation into a single step. Anal. Chem. 2013, 85, 11090–11097.

(22) Kim, S.; Kim, D.; Lee, Y.; Jeon, H.; Lee, B. H.; Jon, S. Conversion of low-affinity peptides to high-affinity peptide binders by using a β-hairpin scaffold-assisted approach. Chembiochem 2015, 16, 43–46.

(23) Landegren, U.; Al-Amin, R. A.; Björkesten, J. A myopic perspective on the future of protein diagnostics. N. Biotechnol. 2018, 45, 14–18.

(24) Zieba, A.; Ponten, F.; Uhlén, M.; Landegren, U. In situ protein detection with enhanced specificity using DNA-conjugated antibodies and proximity ligation. Mod. Pathol. 2018, 31, 253–263.

(25) Xiang, Y.; Nambulli, S.; Xiao, Z.; Liu, H.; Sang, Z.; Duprex, W. P.; Schneidman-Duhovny, D.; Zhang, C.; Shi, Y. Versatile and multivalent nanobodies efficiently neutralize SARS-CoV-2. Science 2020, 370, 1479–1484.

